# Multiplexed imaging of single-synapse activity and astroglial responses in the intact brain

**DOI:** 10.1101/344713

**Authors:** James P. Reynolds, Kaiyu Zheng, Dmitri A. Rusakov

## Abstract

All-optical registration of neuronal and astrocytic activities within the intact mammalian brain has improved significantly with recent advances in optical sensors and biophotonics. However, relating single-synapse release events and local astroglial responses to sensory stimuli in an intact animal has not hitherto been feasible. Here, we present a multiplexed multiphoton excitation imaging approach for assessing the relationship between presynaptic Ca^2+^ entry at thalamocortical axonal boutons and perisynaptic astrocytic Ca^2+^ elevations, induced by whisker stimulation in the barrel cortex of C57BL/6 mice. We find that, unexpectedly, Ca^2+^ elevations in the perisynaptic astrocytic regions consistently precede local presynaptic Ca^2+^ signals during spontaneous brain activity associated with anaesthesia. The methods described here can be adapted to a variety of optical sensors and are compatible with experimental designs that might necessitate repeated sampling of single synapses over a longitudinal behavioural paradigm.

**Highlights:**

- We applied multiplexed multiphoton imaging to optically assess activity of individual synapses and the surrounding astroglia in the intact brain
- Perisynaptic astrocytic regions display localiased, context-dependent Ca^2+^ elevations *in vivo*
- Such elevations may precede ‘spontaneous’ Ca^2+^ entry at presynaptic boutons
- This method paves the way for optical investigation of glutamate release probability *in vivo*

## 1. Introduction

Ca^2+^-dependent, stochastic release of glutamate quanta is a fundamental function of excitatory synapses [1, 2]. The presence of nanoscopic extensions of astrocytes that surround many synapses and demonstrate activity-locked Ca^2+^ excitability has hastened our reevaluation of astrocytic functions and their role in neurotransmission and signal integration in the brain [3–5]. Although various experimental approaches exist for investigating the interplay of perisynaptic astrocyte signals and those of the local synapses, these are technically challenging in organised brain tissue, particularly within the intact brain.

Here, we present an all-optical approach to simultaneous monitoring of Ca^2+^ dynamics within two segmented compartments of the tripartite synapse in response to a physiological stimulus. The technique involves the employment of a multiplexed two-photon microscopy system capable of efficiently exciting spectrally distinct, genetically-encoded optical sensors, which are strategically targeted to various regions of the brain. Thalamocortical projections of the ventrobasal complex (VB) in the thalamus are an ideal target for the selective and sparse labelling of presynaptic elements within the cortex. We demonstrate that labelling of axonal boutons in this way can be coupled with labelling of cortical astrocytes, to reliably yield simultaneous readouts of perisynaptic (astrocytes) and presynaptic (thalamocortical-projecting neurons) Ca^2+^ fluctuations during evoked and spontaneous cortical activity at single synapses. A more exhaustive study of such dynamics might yield important insight into the contribution of astrocytes to different thalamus-mediated behaviours, and indeed the heterogeneity of such contributions [6].

This approach opens up potential routes to assess the impact of local astrocytic Ca^2+^ activity around the synapse, and, once combined with optical glutamate sensor imaging of quantal release as demonstrated earlier [7], should permit live monitoring of neurotransmitter release probability in the intact brain. The latter would be particularly favourable for high-throughput registration of single-synaptic events during learning and behaviour over an extended period.

## 2. Methods

### 2.1 Animals

Animal procedures were carried out under the oversight of the UK Home Office (as per the European Commission Directive 86/609/EEC and the UK Animals (Scientific Procedures) Act, 1986) and in accordance with institutional guidelines. C57BL/6 mice, male and female, were group housed in a controlled environment as mandated by guideline, on a 12 hour light cycle and with food and water provided *ab libitum*.

Genetically-encoded optical sensors were introduced into target CNS cells in young adult mice (p28 - p56) using viral transduction. Cranial windows were subsequently implanted 2 - 3 weeks later and multiphoton microscopy was performed across several imaging sessions, beginning 1 week after the implantation. When performing immunohistochemistry, CNS tissue was dissected from transduced animals, both with and without cranial window implantations, at least four weeks after the viral transduction protocol. Retrograde tracing was carried out in naive wildtype C57BL/6 mice. Details for each of these methods are given below.

### 2.2 AAV transduction

C57BL/6 mice were prepared for aseptic surgery and anaesthetised using isoflurane (5% v / v induction, 1.5 - 2.5% maintenance). The scalp was shaved and disinfected using chlorhexidine, topically applied in three sequential washes. The animal was secured in a stereotaxic frame (David Kopf Instruments, CA, USA) and the appropriate depth of anaesthesia was confirmed before continuing. Body temperature was maintained at 37.0 ± 0.5 °C using feedback control. Perioperative care included analgesia (subcutaneous buprenorphine, 60 μg kg^−1^, topical lidocaine / prilocaine emulsion, 2.5% / 2.5%) and ocular protection (Lacri-lube, Allergan, UK). The scalp and underlying superficial tissue were parted at the midline and the skull was exposed. Two craniotomies of approximately 1 - 2 mm diameter were carried out using a hand drill (Proxxon, Föhren, Germany) at sites overlying the thalamic VB and / or the somatosensory cortex (S1). When combined, the injection into the VB was completed prior to exposing S1. During VB transduction, stereotactic coordinates injections were adjusted to favour the ventral posteromedial nucleus (VPM) or the ventral posterolateral nucleus (VPL) and were as follows; VPM, 1.8 mm posterior, 1.5 mm lateral and 3.2 mm dorsal, relative to bregma; VPL, 1.6 mm posterior, 1.75 mm lateral and 3.2mm dorsal, relative to bregma. The total injection volume was delivered in three steps, reducing dorsoventral depth by 100 μm at each step. For S1 injections, a single bolus was delivered at a depth of 600 μm. The coordinates when targeting the barrel cortex were 0.5 mm posterior and 3.0 mm lateral to bregma. The coordinates for the S1 forelimb region were 0.2 mm posterior and 2.0 mm lateral to bregma. A warmed saline solution was applied to exposed cortical surface throughout the procedure.

Pressure injections of adeno-associated virus (AAVs, totalling between 0.2 and 1 × 10^10^ genomic copies, in a volume not exceeding 500 nL, supplied by Penn Vector Core, PA, USA) were carried out using a glass micropipette, at a rate of ~1 nL sec^−1^, stereotactically guided to the target regions as outlined above. AAVs using in this study comprised; AAV9.Syn.GCaMP6f [8]; AAV1.Syn.NES-jRCaMP1b [9]; AAV5.GfaABC1D.Lck-GCaMP6f [10]; and AAV5.GfaABC1D.cyto-tdTomato. Once delivery was completed, pipettes were left in place for 5 minutes before being retracted. The surgical wound was closed with absorbable 7-0 sutures (Ethicon Endo-Surgery GmbH, Norderstedt, Germany) and the animal was monitored and recovered in a heated chamber. Postoperative analgesia (meloxicam, subcutaneous, 1 mg kg^−1^) was administered at least once daily for up to two days following surgery. Mice were then prepared for cranial window implantation approximately 2 - 3 weeks later.

### 2.3 Cranial Window Implantation

Mice were prepared for aseptic surgery and secured in a stereotaxic frame as above. A large portion of the scalp was removed to expose the right frontal and parietal bones of the skull, as well as the medial aspects of the left frontal and parietal bones. To facilitate cement bonding during fixation of the cranial window implant, the right temporalis muscles were retracted laterally to expose the squamous suture. The skull was coated with Vetbond (3M, MN, USA) and a custom-made titanium headplate was placed over the target region of S1. The assembly was further secured with dental cement (SuperBond, Sun Medical Co. Ltd., Japan). The animal was then removed from the stereotaxic frame and secured, via the introduced headplate, in a custom-built head fixation frame. A craniotomy with a diameter of 4 mm at the outer edges was carried out over the right somatosensory cortex, centred over the injection site in S1. Immediately prior to removal of the skull flap, the surface was superfused with warmed aCSF (in mM; 125 NaCl, 2.5 KCl, 26 NaHCO_3_, 1.25 Na_2_HPO_4_,18 Glucose, 2 CaCl_2_, 2 MgSO_4_; saturated with 95% O_2_ / 5% CO_2_, pH 7.4). The exposed dura was carefully excised using a combination of 26G needles (pushed against a hard surface to introduce a curved profile), fine-tipped forceps (11252-40, Fine Science Tools, Germany) and 2.5 mm spring scissors (15000-08, Fine Science Tools, Germany), taking care not to penetrate to the pia mater. Next, a previously prepared coverslip consisting of a 3 mm diameter round coverglass affixed beneath a 4 mm diameter round coverglass (Harvard Apparatus UK, bonded using a UV-curable optical adhesive (NOA61), ThorLabs Inc., NJ, USA) was placed over the exposed cortex. Downward pressure was applied to the coverslip using a guided wooden spatula. To prevent excessive force, the spatula was previously severed and sanded to allow flexibility. Superfusion was discontinued and excess fluid was removed using a sterile surgical sponge, taking care not to wick fluid from beneath the cranial window. VetBond was carefully applied around the edges of the cranial window to seal and secure the coverslip. The wooden spatula was raised and a final layer of dental cement was applied. Once all bonding agents had cured, the animal was recovered in a heated chamber and returned to its homecage when ambulatory. Post-operative care was administered as before during the viral transduction procedure.

### 2.4 Multiplexed multiphoton microscopy

Two-photon microscopy was performed to simultaneously capture relative subcellular changes in Ca^2+^ Genetically-encoded optical sensors were used to target distinct populations, which were then sampled from using a wavelength multiplexing suite comprised of a Newport-Spectraphysics Ti:sapphire MaiTai tunable IR laser pulsing at 80 MHz and a Newport-Spectraphysics HighQ-2 fixed-wavelength IR laser pulsing at 63 MHz. The laser lightpaths were aligned (though not synchronised) before being point-scanned using an Olympus FV1000 with XLPlan N 25x water immersion objective (NA 1.05). Animals were lightly anaesthetised with an injectable triple-anaesthetic mix (fentanyl, 0.03 mg kg^−1^, midazolam, 3 mg kg^−1^, and medetomidine, 0.3 mg kg^−1^) and secured under the objective on a custom-built stage, via the implanted titanium headplate. An additional dose (fentanyl, 0.015 mg kg^−1^, midazolam, 1.5 mg kg^−1^, and medetomidine, 0.15 mg kg^−1^) was applied approximately every 90 minutes thereafter.

Initially, to locate labelled thalamocortical boutons in S1 within the arbor of labelled cortical astrocytes, z-stacks of high-spatial resolution were acquired with both lasers illuminating the tissue at 910 nm (to excite green indicators) and 1040 nm (to excite RCaMP1b), favouring slow acquisition rates and long-dwell times to visualise subcellular structures. Z-stacks were 512 × 512 pixels, with a pixel size of 0.25 - 0.5 μm and an interval size of 1 - 5 μm. Subsequently, timelapse measurements of higher temporal resolution were performed in L1 and L2/3, at depths of 50 - 150 nm. For bouton recordings, framescans of 4 - 20 Hz were performed, with a pixel dwell time of 2 μs. Resolutions and pixel size varied by recording. Mean laser power at the focal plane was measured at 20 - 50 mW and adjusted according to depth and sampling rates. To verify that boutons were active and involved in sensory information processing, brief 5 second, 3 Hz stimulation protocols were carried out. For S1 barrel field, air puffs of nitrogen were directed at the contralateral whiskers. For imaging within the forelimb region, an isolated constant current stimulator was coupled to a subcutaneous bipolar electrode and an electrical stimulus (0.2 mA) was delivered to the contralateral forelimb, across the third interdigital pad. Precise synchronisation and triggering was controlled by Spike2 (Cambridge Electronic Design Ltd., UK, v2.7) and physiological stimulations were delivered using the 1401 laboratory interface (Cambridge Electronic Design Ltd., UK). All recordings were coded (for blinding where necessary) and motion-corrected using MATLAB. Custom-written MATLAB scripts were used to segment timelapse images and define the relative change in Ca^2+^-dependent fluorescence.

### 2.5 Retrograde Tracing

Naive C57BL/6 mice (p30 - 50) were anaesthetised and prepared for aseptic intracranial injection as outlined above for viral transduction. A 1 - 2 mm craniotomy was carried out and FluoroGold (4% in distilled water) was pressure-injected into layer IV of S1 using a pulled-glass micropipette in order to retrogradely label S1-projecting neurons in the thalamus. The animals were recovered and killed 7 - 10 days later by transcardial perfusion with saline and 4% formaldehyde solution. The brain was removed and immersed in formaldehyde solution for 24 hours. After washing in PBS, the tissue was sectioned (50 μm coronal sections, collected using a Leica VT1000S vibratome) and imaged using epifluorescent microscopy.

### 2.6 Immunofluorescence

After viral transduction or imaging experiments, whole brain tissue was dissected following transcardial perfusion with saline followed by transcardial fixation with formaldehyde solution (4%). Brains were left immersed in formaldehyde solution for 24 hours and stored in PBS at 4°C until sectioning. 30 μm sections were collected using a Leica VT1000S vibratome and were stored at −20°C in cryoprotective medium (30% glycerol and 30% ethylene glycol in 1X PBS). In preparation for immunolabelling, sections were washed (PBS, 3 × 5 min), permeabilised (0.1% Triton-X in PBS, 15 min), washed in glycine (1M in PBS, 30 min) and blocked (1% bovine serum albumin in PBS, 60 min). Sections were incubated in blocking buffer overnight at 4°C, along with the following primary antibodies; anti-GFP (1:800, Thermo Fisher Scientific #A10262), anti-RFP (1:400, Rockland Immunochemicals Inc. #600-401-379, anti-VGLUT1 (1:1000, Merck-Millipore #AB5905). After washing, immunoreactivity was visualised using AlexaFluor 488, 568 and 594 secondary antibodies (1:500 dilution in blocking buffer, Thermo Fisher Scientific #A11075, #A11039, #A11020 and #A11072), incubating at room temperature for 2 hours. Tissue was then washed and mounted in Vectashield hard setting mounting medium (Vector Laboratories Ltd., UK), with or without DAPI nuclear staining. Images were captured an epifluorescence microscope.

### 2.7 Statistics

To assess differences in astrocytic Ca^2+^-dependent fluorescence levels before, during and after defined presynaptic Ca^2+^ transients (Fig. 2i), mean ΔF/F values for the corresponding periods were compared by Mann-Whitney U-tests, and significance was accepted at ρ < 0.05. For spontaneous presynaptic Ca^2+^ entry events, the ‘before peak’ values were taken from the period immediately before the transient elevation in presynaptic Ca^2+^, while for the evoked responses, the ‘before peak’ values were taken immediately before the onset of the stimulation protocol. ‘After peak’ values correspond to a period of 1.5 - 4.5 seconds after the peak presynaptic Ca^2+^ level.

## 3. Results

### 3.1 Dual AAV transduction of the ventrobasal complex and somatosensory cortex effectively labels thalamocortical boutons within astrocytic arbors

Astrocytes are ubiquitous among all cortical layers [11] and are sensitive to neurotransmission through ultrathin perisynaptic processes that approach and surround synapses [12–14]. In order to investigate astrocyte-neuron networking in the live animal, we envisaged an experimental setup in which indicators of both astrocytic and synaptic activity were measured through dual-population multiphoton microscopy. Due to the density and heterogeneity of cortical synapses, we sought to selectively and sparsely label presynaptic boutons originating from a distant nucleus of the brain. We first employed retrograde tracing to identify thalamic nuclei involved in the relaying of sensory information to S1. FluoroGold was injected into the whisker- and forelimb-associated regions of S1 in C57BL/6 mice, and histological sections were taken 7 days later. As previously described [15], retrograde tracing revealed that two distinct nuclei of the VB in the thalamus project to S1 whisker- (S1BF) and S1 forelimb-(S1FL) associated regions of S1; the VPM and the VPL, respectively (Fig. 1a).

**Figure 1:**
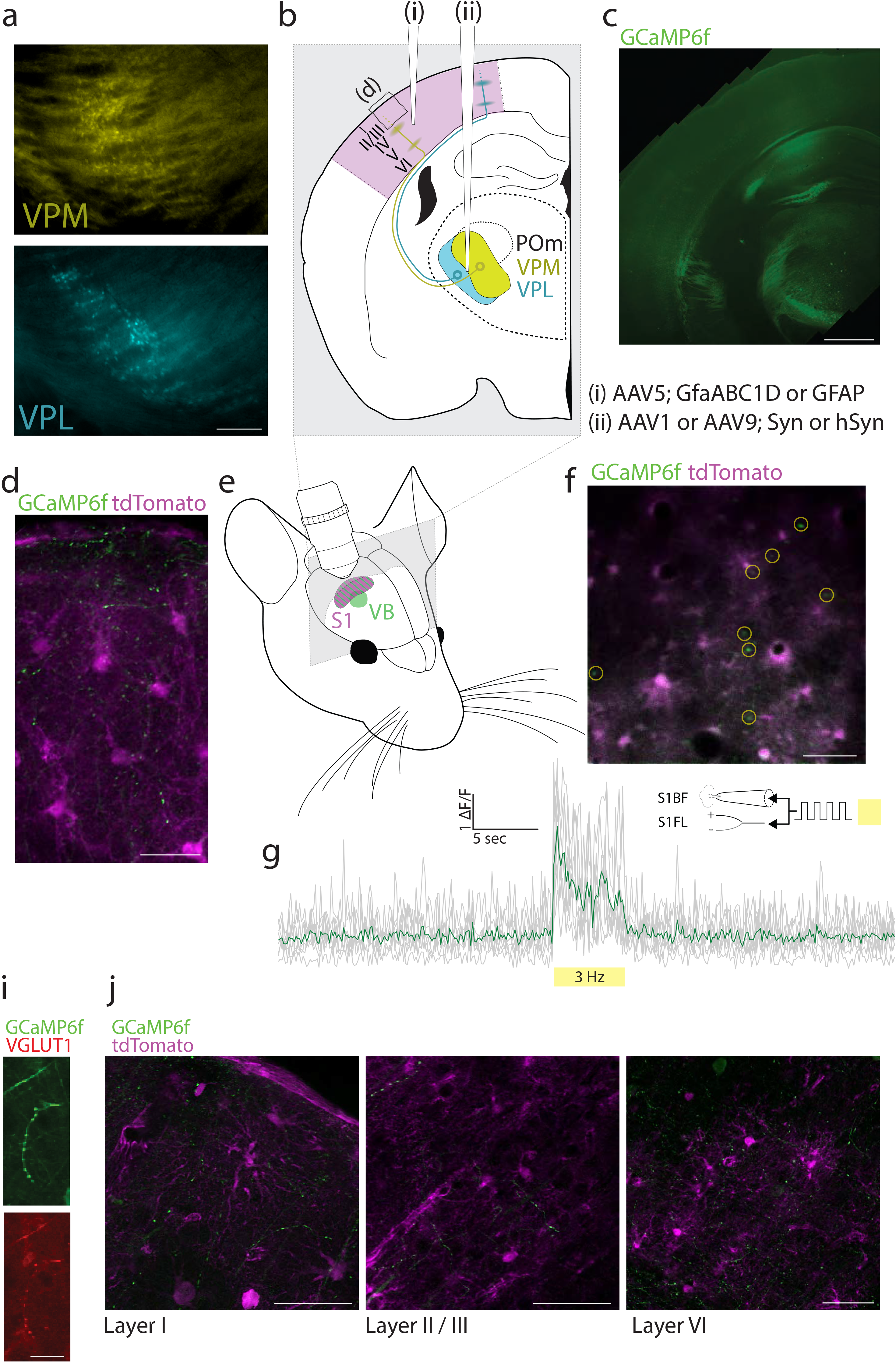
Experimental approach to labelling axonal boutons within the cortical astrocyte arbor. **a,** Regions of the ventrobasal complex of the thalamus identified through retrograde tracing following injection of FluoroGold into the whisker- (top, labelling ventral posteromedial thalamus, VPM) and forepaw-associated regions (bottom, labelling ventral posterolateral thalamus, VPL) of the somatosensory cortex. Scale, 200 μm. **b,** AAVs were injected into the somatosensory cortex (i) and ventrobasal complex (ii) to label trespassing axons within the arbor of cortical astrocytes. **c,** VPL injection of viral constructs expressing GCaMP6f. Scale, 1 mm. **d,** confocal images of immunolabelled fixed cortex (Layer 1, as indicated in **b**, boxed region) following AAV injections of the cortex and the ventrobasal complex, as outlined in **b**. Scale, 30 μm. **e,** Experimental setup outlining the targeted regions for labelling the somatosensory cortical boutons and astrocytes, and subsequent multiphoton imaging. **f, g** Multiphoton imaging of GCaMP6f-positive thalamocortical boutons in the whisker region of somatosensory cortex during tactile stimulation, where astrocytes are also labelled (tdTomato). **f** is an image projection (layer II / III, depth 100 μm) over the duration of the stimulus and **g** is the Ca^2+^-dependent fluorescence in the highlighted boutons (**f**, yellow ROIs). Scale, 50 μm. **i,j** Confocal images of immunolabelled fixed cortex. VGLUT1-positive boutons were observed throughout the cortical layers. Scales, 15 μm (**i**) and 50 μm (**j**).

**Figure 2:**
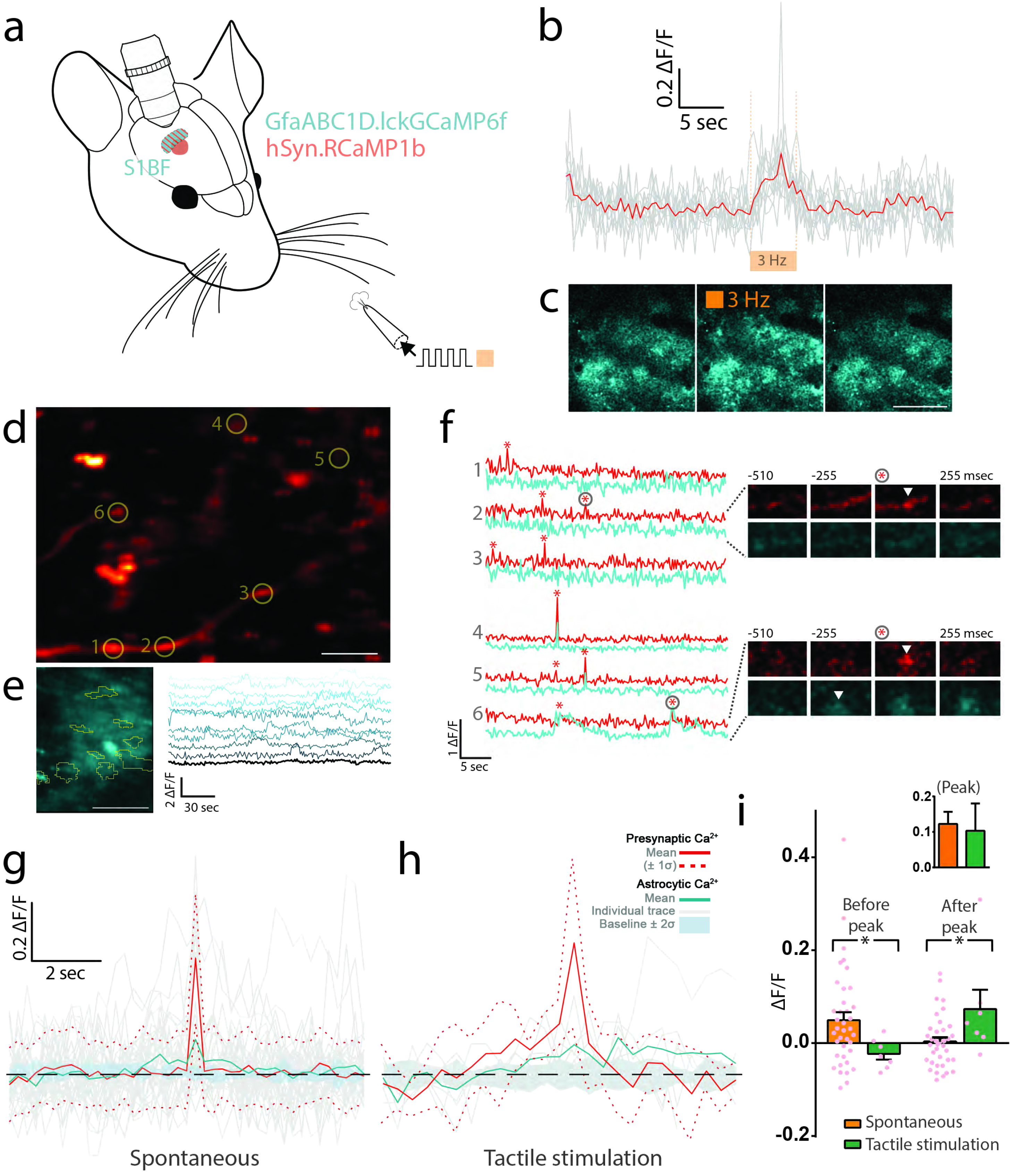
Investigating the heterogeneity of astrocytic calcium transients during synaptic transmission. **a**, Experimental setup for labelling of the somatosensory cortical boutons and astrocytes with the indicated viral constructs. **b**, Ca^2+^-influx within labelled boutons during tactile stimulation of the whiskers. **c**, Synchronous Ca^2+^ propagations throughout multiple astrocytes during tactile stimulation as in **b**. Scale, 150 μm **d**, The whisker cortical region (depth 40 μm) showing RCaMP1b expressed in the axons of thalamocortical neurons following AAV injection into the ventral posteromedial thalamus; circles, ROIs. Scale, 10 μm. e, Spontaneous Ca^2+^ transients within the arbor of a single astrocyte. Scale, 20 μm. **f**, Registration of Ca^2+^ fluctuations within segmented compartments of a single synapse, from ROIs in **d**. Perisynaptic Ca^2+^transients (membrane-tethered GCaMP6f, cyan) and presynaptic Ca^2+^ influx (RCaMP1b, red) were observed during spontaneous activity in an anaesthetised brain; * indicates 4σ, four standard deviations above the baseline noise. Timelapse montages are shown for a single event within two synapses (ROIs 2 and 6 as in **d**). **g, h,** Presynaptic Ca^2+^-dependent fluorescence (red trace, dashed lines represent ± standard deviation) was aligned at the peak fluorescence values for a given synaptic event, and the corresponding perisynaptic astrocytic Ca^2+^ levels were plotted. Individual (gray) and mean (cyan) traces are shown. A ‘baseline’ astrocytic Ca^2+^ trace was also computed by sampling astrocytic fluorescence data at randomised timepoints during imaging acquisitions that did not involved tactile stimulation (thick cyan trace representing the computed baseline value ± 2 standard deviations). **i**, ΔF/F values were computed for specified periods before, during and after peak presynaptic Ca^2+^ levels as in **g**, **h**

Next, we targeted both VB and S1 with AAVs expressing genetically-encoded fluorescent proteins in order to assess the possibility of measuring presynaptic activity within the territory of labelled cortical astrocytes (Fig. 1b). AAVs targeted at cortical astrocytes in S1 contained a GFAP or GFAP-based promoter sequence, while those targeted the VB contained a synapsin promoter. We transduced VB neurons with the genetically-encoded Ca^2+^ indicator (GECI) GCaMP6f, and S1 astrocytes with the red fluorescent protein tdTomato. Immunohistochemistry revealed effective transduction of VB neurons as evident by dense somatic labelling within the VB and visible projections within white matter tracts of the caudate putamen, the internal capsule, and the external capsule (Fig. 1c). We performed confocal imaging of such sections to investigate cortical staining patterns in better detail, demonstrating that GCaMP6f-positive varicosities resembling boutons are visible throughout the cortex, even within superficial layers (Fig. 1d). These structures are positive for vesicular glutamate transporter 1 (VGLUT1, Fig. 1i) and are most dense at deeper layers (IV, V, and VI), in keeping with previous descriptions of the canonical thalamic projections (Fig. 1j, see [16]). Axonal ramifications were nonetheless observed at lower densities within layers I - III (Fig. 1j), as was previously reported for thalamocortical projections to somatosensory [17] and visual cortex [18]. Importantly, within targeted regions of S1, axonal projections trespassed the arbor of tdTomato-positive astrocytes across all layers (Fig. 1d, 1j).

Next, we confirmed that tactile stimulation evoked rapid Ca^2+^ elevations in GCaMP6f-positive boutons. Two-four weeks after AAV transduction, a cranial window was implanted over S1BF or S1FL(Fig. 1e). Once recovered (> 1 week after implantation), multiplexed multiphoton imaging was employed to capture both tdTomato-positive astrocytes and GCaMP6f-positive boutons in S1, in anaesthetised mice (Fig. 1f). During a brief tactile stimulation protocol (5 sec, 3 Hz) involving either air puffs (S1BF) or subcutaneous electrodes (S1FL), rapid elevations of Ca^2+^ were observed during the stimulation period across a large number of segmented boutons, indicative of presynaptic Ca^2+^ entry during action potential firing of VB neurons and as illustrated here in S1BF following contralateral whisker puffs (Fig. 1g).

### 3.2 Perisynaptic astrocytic calcium transients coincide with spontaneous and evoked calcium entry at thalamocortical boutons *in vivo*

Having established a method for simultaneous imaging of presynaptic and astrocytic compartments in C57BL/6 mice *in vivo*, we next leveraged a membrane-tethered Ca^2+^ indicator, lck-GCaMP6f [10], to record spectrally separate Ca^2+^ transients within both the ultrathin astrocytic processes and their neighbouring boutons labelled with a red-shifted Ca^2+^ indicator, RCaMP1b (Fig. 2a). AAV injections were targeted to S1BF and VPM, respectively. Though RCaMP1b is dimmer than GCaMP6f, with reduced dynamic range [9], we continued to observe robust Ca^2+^ elevations during brief, tactile stimulation of the contralateral whiskers (Fig. 2b) during anaesthesia. Evoked presynaptic Ca^2+^ elevations were accompanied by highly synchronous Ca^2+^ elevations across numerous astrocytes (Fig. 2c). Outside of tactile stimulation, thalamocortical boutons in S1 layer II / III were generally quiescent, but displayed ‘spontaneous’ low-frequency, short-lived (< 250ms) elevations of Ca^2+^-dependent fluorescence (Fig. 2d, 2f), while astrocytes displayed localised, stochastic Ca^2+^ transients that lacked significant network coherence often seen during awake behaviours [19, 20], as seen in Fig. 2e. Although such bouton activity was not always accompanied by astrocytic Ca^2+^ transients (see Fig. 2f, top panel, ROIs 1-3), we observed some correlated Ca^2+^ propagations within astrocytes during spontaneous Ca^2+^ entry at boutons (see Fig. 2f, bottom panel, ROIs 4-6).

In order to generate a better measure of the dynamics of these co-occurrences, we time-matched the peaks of Ca^2+^-dependent fluorescence within RCaMP1b-positive boutons and assessed the perisynaptic astrocytic Ca^2+^ signal. Peaks were defined as local maxima when fluorescence within boutons exceeded a value greater than 4 standard deviations above the mean fluorescence. At peak bouton Ca^2+^-dependent fluorescence, Ca^2+^ levels within the overlapping perisynaptic astrocytic regions increased during both spontaneous and stimulation-evoked events, with no difference noted in the intensity of the signal (Fig. 2i, inset). Surprisingly, however, we found that perisynaptic astrocytic Ca^2+^ levels increased above baseline levels prior to spontaneous Ca^2+^ entry at boutons (Fig. 2g). During stimulation-evoked presynaptic Ca^2+^ elevations, the astrocytic Ca^2+^ signal lags behind that of the boutons but remains elevated after the cessation of stimulation (Fig. 2h). Directly comparing these dynamics revealed a significant divergence in the timing of astrocytic Ca^2+^ transients within the perisynaptic regions, dependent of the nature of the Ca^2+^ signal within boutons (*ρ < 0.05, Mann-Whitney U-test, Fig. 2i).

## 4. Discussion

In the present study, we demonstrated a simple viral labelling approach to target and simultaneously image two constituent components of the thalamocortical synapse, successfully implementing this strategy to advance an experimental method for optical monitoring of individual synaptic events *in vivo*.

The approach involved pervasive labelling of entire astrocytic arbors and sparse labelling of presynaptic boutons along thalamocortical projections *in vivo*, using spectrally-separable Ca^2+^ indicators. Recent advances in imaging acquisition technologies and genetically-engineered optical sensors are facilitating such approaches in relating synaptic transmission to the activity of nearby perisynaptic astrocytic processes. In acute slices, single-pulse stimuli applied to the perforant path are sufficient to reliably evoke time-correlated, localised Ca^2+^ transients within fine astrocytic processes proximal to the imaged axonal projections which represent some of the stimulated fibers [21]. Evoked astrocytic Ca^2+^ transients such as these are sensitive to TTX, modulated by synaptically released factors including glutamate and ATP [22] and partially dependent on intracellular stores within the endoplasmic reticulum [5, 23] (although see [24]). However, other modalities of astrocytic Ca^2+^ signals that do not depend on excitatory transmission have been reported; these may be stochastic, TTX-insensitive events [21], dependent on extracellular Ca^2+^ [25, 26], or responsive to purinergic [27–29], domapinergic [30], or GABAergic [31, 32] signals, in addition to other wide-ranging ‘homeostatic’ influences such as those of mechanical of vascular origin [33]. A range of potential Ca^2+^ sources for a given propagation of intracellular astrocytic Ca^2+^ have been proffered and likely represent distinct astrocytic compartments and network activity contexts [34–36].

We thus achieve simultaneous pre- and perisynaptic astroglial Ca^2+^ monitoring in the mammalian brain during physiologically evoked or spontaneous synaptic activity, observing divergent patterns of activity depending on the nature of synaptic discharges. Tactile stimulation evoked both axonal burst activity (manifest here as a transient elevation in presynaptic Ca^2+^ levels throughout the stimulation) and synchronous astrocytic Ca^2+^ transients throughout the neuropil. Though elevations of astrocytic Ca^2+^ within the perisynaptic region are faster than the classic somatic astrocytic Ca^2+^ transients, they lag behind the rapid onset of synaptic transmission, presumably responding to continued glutamate release onto perisynaptic astrocytic membranes. However, these Ca^2+^ elevations persist beyond the cessation of tactile stimulation, lasting several seconds after presynaptic Ca^2+^ levels have returned to baseline. These observations may reflect the notion of temporal (and possibly spatial) signal integration represented by astroglial Ca^2+^ activity in response to intermittent synaptic discharges [4, 34].

Our proof-of-principle data unveil a surprising relationship between perisynaptic astrocytic Ca^2+^ transients and spontaneous Ca^2+^ entry at boutons, whereby in the absence of tactile stimulation, low-frequency stochastic elevations in presynaptic Ca^2+^ are preceded by astrocytic Ca^2+^ in the perisynaptic region. Whilst the exact nature of this relationship needs a dedicated study, it is likely that local astrocyte activity alone could not be the source for presynaptic spike-dependent Ca^2+^ increases as action potentials in this context are generated distally. Persistent thalamocortical loops involved in tactile discrimination and motor planning [37] may induce activity both locally (i.e., in cortical astrocytes) and distally (in the thalamocortical projecting neurons). In certain conditions, receptor-triggered Ca^2+^ release from Ca^2+^ stores in axonal boutons could elevate presynaptic Ca^2+^, although such events are likely to be rare [38]. The present study does not capture data to satisfactorily assess the number and frequency of action potentials occurring during the observed spontaneous Ca^2+^-influx into boutons. However, relative to the observed synchrony during tactile stimulation (see Fig. 1f and 1g), astrocytic modulation of spontaneous Ca^2+^-influx is plausible and may be worth investigating further.

One key advantage in employing viral strategies is the flexibility in targeting different brain regions and cell types. Targeting can be refined using genetic control, indicible elements, transgenic animals and other such strategies to yield functionally relevant synaptic couples. For instance, layer-specific transduction of single barrels has been demonstrated [39], and it is feasible that individual cells be targeted (and characterised *post-hoc*) by electroporation [40] at the time of cranial window implantation. Coupled with the targeting of thalamocortical boutons, it might be possible to selectivity label presynaptic boutons and postsynaptic dendritic spines of defined neural populations with any number of desired optical sensors. For instance, this method offers an all-optical approach to measuring glutamate release probability *in vivo* if combined with an optical glutamate sensor to image quantal release, as previously demonstrated [7]. This approach would readily permit repeated sampling of the same synaptic release site during longitudinal experiments assessing learning and behaviour.

## Acknowledgements

We would like to acknowledge the GENIE Program and the Janelia Research Campus for the provision of materials to Penn Vector Core which were used in this study; namely, vectors encoding GCaMP6f and RCaMP1b. We specifically thank Vivek Jayaraman, Ph.D., Douglas S.Kim, Ph.D., Loren L. Looger, Ph.D., and Karel Svoboda, Ph.D. from the GENIE Project, Janelia Research Campus, Howard Hughes Medical Institute. Similarly, we would like to thank Bhaljit Khakh, Ph.D. from the University of California, Los Angeles, for providing the construct encoding the astrocyte-specific lck.GCaMP6f sensor.

## Author Contributions

J.P.R. designed and performed experiments; K.Z. advised on optical designs; D.A.R. conceived the study; All authors contributed to the manuscript.

## Declaration

The authors declare that there are no competing financial or personal interests.

